# DeMAG predicts the effects of variants in clinically actionable genes by integrating structural and evolutionary epistatic features

**DOI:** 10.1101/2022.06.15.496230

**Authors:** Federica Luppino, Ivan A. Adzhubei, Christopher A. Cassa, Agnes Toth-Petroczy

**Affiliations:** Max Planck Institute of Molecular Cell Biology and Genetics, Dresden 01307, Germany; Center for Systems Biology, Dresden 01307, Germany; Cluster of Excellence Physics of Life, TU Dresden, 01062 Dresden, Germany; Brigham and Women’s Hospital Division of Genetics, Harvard Medical School, Boston, MA, 02115 USA; Department of Biomedical Informatics, Harvard Medical School, Boston, MA 02115 USA

## Abstract

Despite an increasing use of genomic sequencing in clinical practice, interpretation of rare genetic variants remains challenging even in well-studied disease genes, resulting in many patients with Variants of Uncertain Significance (VUSs). Computational Variant Effect Predictors (VEPs) are currently used to provide valuable evidence in variant classifications, but they often misclassify benign variants, contributing to potential misdiagnoses. Here, we developed Deciphering Mutations in Actionable Genes (DeMAG), a supervised classifier for interpreting missense variants in actionable disease genes with improved performance over existing VEPs (20% decrease of false positive rate). Our tool has balanced specificity (82%) and sensitivity (94%) on clinical data, and the lowest misclassification rate on putatively benign variants among evaluated tools. DeMAG takes advantage of a novel epistatic feature, the *‘*partners score’, which is based on evolutionary and structural partnerships of residues as estimated by evolutionary information and AlphaFold2 structural models. The ‘partners score’ as a general framework of epistatic interactions, can integrate not only clinical but functional information. We anticipate that our tool (demag.org) will facilitate the interpretation of variants and improve clinical decision-making.

## Introduction

Assessing the pathogenicity of genetic variants remains a significant challenge in research and clinical translation. The American College of Medical Genetics and Genomics (ACMG) recommends reporting secondary findings which are known to be pathogenic in a set of clinically actionable genes (e.g., ACMG SF lists^1,2^) for patients who undergo sequencing^3^. Knowledge of a pathogenic variant in such a gene might improve clinical management, diagnosis, and prevention. At present, over three quarters of variants which have been submitted to ClinVar^4^ are classified as Variants of Uncertain Significance (VUSs) given insufficient epidemiological, functional, or other supportive evidence (Supplementary Fig. 1). Importantly, many patients who carry variants in these established disease genes will not learn about them when following clinical guidelines, despite their potential for increasing risk of disease. The uncertainty about the pathogenicity of a variant may pose a psychological burden for patients, left without guidance, and can lead to potential health costs associated with under and overdiagnosis.

Many Variant Effect Predictors (VEPs) have been developed to predict the functional impacts of these variants, and these tools are often used in diagnostic variant interpretation. A computational prediction that a variant will have a deleterious effect is considered evidence in support of pathogenicity when following the American College of Medical Genetics and Genomics/Association for Molecular Pathology (ACMG/AMP) clinical guidelines for sequence variant interpretation^5^. Widely used predictors include PolyPhen-2^6^, VEST4^7^, M-CAP^8^, and REVEL^9^. Except REVEL and VARITY^10^ which are meta-predictors, the others are supervised methods which use lists of pathogenic and benign variants to train a statistical model that assigns a pathogenicity score for any given variant using sequence-based and structural features. While most tools are designed to be used exome-wide, specialized predictors can reach higher performance on selected genes and disease phenotypes^11^.

Unsupervised methods, such as DeepSequence^12^, EVmutation^13^ and EVE^14^ are agnostic to variant labels as they infer functional effects from multiple sequence alignment (MSA) alone. These methods rely on the availability of sufficient quality MSA data, which is often missing in disordered and low-complexity regions, and poorly conserved genes. Unsupervised methods characterize the fitness effects of mutations independently from reported disease-causing variants, and do not provide an interpretation of pathogenicity^13,15^, with the exception of EVE, which relies on labeled clinical data to identify gene-specific pathogenicity thresholds. While this is useful for clinical applications, it suffers from the same labeling bias as other supervised tools.

Due to limited clinical data, there are two primary challenges in training sufficiently accurate VEPs^16^. The first issue (type 1 circularity) refers to a biased testing set and requires that the testing set contains variants that were not used in the training of all predictors. This is challenging as many methods train models using variants collected from similar sources, and can result in general inflation of predictive performance. The second issue (type 2 circularity) refers to an intrinsic characteristic of clinical databases: variants in a given gene, with an established link to a disease phenotype, may often be classified as pathogenic^17^. VEPs which use gene-based features, e.g., length of the protein, can make predictions based on a gene’s characteristics and pathogenicity, rather than on the attributes of a specific variant. This bias hinders discrimination between pathogenic and benign variants within a given gene and skews the predictive performance toward high sensitivity and poor specificity^18^. Thus, addressing these issues of unbiased testing set, and balancing sensitivity and specificity are crucial for the development of an accurate predictor for clinical applications.

Evolutionary sequences and 3D structures of proteins contain valuable information about the importance of residue positions and substitutions. The evolutionary conservation of a position in orthologous sequences correlates with the tolerance to mutations within a population, and can be used to predict the pathogenicity of genetic variants^19^. Several conservation scores have been developed and are used as predictive features in VEPs^19–21^. While most assume site-independence, considering epistasis between pairs of residue positions improves variant assessment^13^. Here, epistasis refers to the interdependence of two residue positions. An estimated 90% of variation is impacted by epistasis^22,23^. Indeed, human pathogenic variations appear as neutral substitutions in closely-related orthologous from other species. These are termed, compensated pathogenic deviations^24^ as the pathogenicity of the substitution is suppressed by another compensatory substitution either within the same gene^25–28^ or in another one^29^. The compensatory mechanism often involves residues in close proximity in the 3D structure and the preservation of side-chain side-chain interactions^24^. In general, the hydrophobic core of proteins tends to evolve slowly, while the surface evolves faster^30^. Accordingly, disease-causing mutations tend to occur in the hydrophobic core of the 3D structure of the protein, while common variants tend to be located on the surface, i.e. areas with high solvent accessibility^31^. Incorporating epistatic and structural information comprehensively only recently became possible with biome-wide sequencing efforts and AlphaFold2 predictions^32^.

Here, we extend the traditional conservation paradigm to assess variant effects with novel protein sequence-and structure-based features. We designed an epistatic feature, the partners score, which defines epistatic residue pairs based on co-evolutionary and 3D structural partnership of residues as defined by AlphaFold2^32^ models. The partners score is informed by the clinical label of the partner residues, and it relies on the wealth of already existing clinical knowledge. Based on their high medical relevance and wealth of clinical diagnostic data, we focused on interpreting missense variants in clinically actionable disease genes in the ACMG SF v2.0 list, which we refer to as ACMG SF genes^2^.

We developed DeMAG (Deciphering Mutations in Actionable Genes), a specialized supervised classifier for the ACMG SF genes. DeMAG achieves the best performance across VEPs tested on clinical and common variants in population sequencing data. Our specialized classifier outperforms VEPs not designed for ACMG SF genes, which are critical for clinical sequencing applications. We share predictions and interpretations of all ∼1.3 million missense variants in the ACMG SF genes as a web server application (demag.org).

## Results

### Method overview

The uncertainty associated with the interpretation of mutations in clinically actionable disease genes presents major challenges for clinical translational research. Under and overdiagnosis as well as patients’ psychological burden due to lack of evidence to support variants’ pathogenicity may result in increased costs for the healthcare system. Therefore, we developed DeMAG (Deciphering Mutations in Actionable Genes) a supervised classifier to assess the pathogenicity of mutations in clinically actionable disease genes (ACMG SF) and support clinical decision making. First, we carefully curated variants with known phenotypic effects and putatively benign variants used for training the model (Fig. 1 and Supplementary Fig. 2). For those variants, we then tested several sequence-and structure-based features and selected those that discriminated between pathogenic and benign mutations (Fig. 1 and Supplementary Table 1). We designed the partners score, which is based on evolutionary and structural partnerships of residues as estimated by AlphaFold2 structural models. Then, we trained a machine learning model that was validated with 3 different ground-truth test sets: clinical, functional (deep mutational scanning) data, and common variants from population data. Finally, we computed DeMAG pathogenicity scores for all missense variants in the ACMG SF genes.

**Figure 1:**
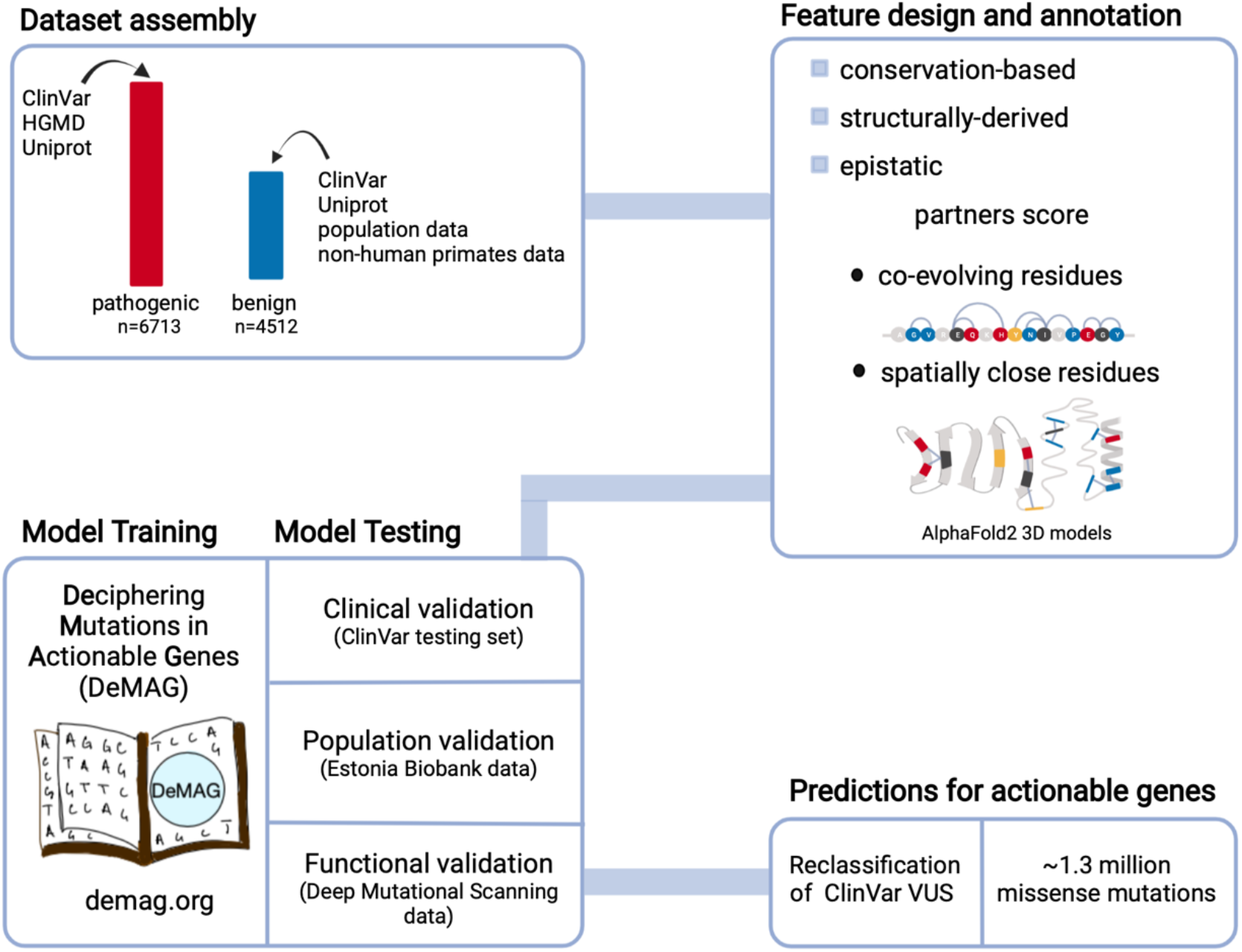
DeMAG (Deciphering Mutations in Actionable Genes) overview. First, we assembled the training set for 59 actionable genes; the pathogenic variants were collected from clinical databases such as ClinVar and HGMD. The benign class was enriched with variants from different sources such as clinical, population and benign human orthologues data. The training set consisted of 6713 (60%) pathogenic and 4512 (40%) benign variants. Next, we annotated the features, e.g., EVmutation, IUPred2A and AlphaFold2 confidence score pLDDT. We designed a novel feature, the partners score feature, that captures the role of epistasis both in the sequence and in 3D space of the protein. It highlights how evolutionary coupled and spatially close residues are enriched in the same phenotypic effect. Next, we trained a classification tree based gradient boosting model using the selected 13 features. The model was validated with 3 different types of data, 1) clinical testing set from ClinVar, 2) putatively benign common variants from the Estonian population, 3) Deep Mutational Scanning (DMS) data for four genes, i.e., *BRCA1, TP53, MSH2* and *PTEN*. The validation sets identify variants with a ground-truth label, e.g., variants’ clinical significance or functional score. Finally, we provide predictions for all variants whose phenotypic effect is yet unclear or unknown.

### Curated training set

In order to curate a high-quality training set, we considered several independent sources of SNVs and set strict criteria to retain only high-quality variants. We included high-quality ClinVar variants with at least two stars review status label, namely variants labeled with no conflicts between all submitters or reviewed by expert panels or practice guidelines. We supplemented that with variants which have previously been described in the medical literature in Human Gene Mutation Database (HGMD)^33^, that have not yet been observed in ClinVar (Supplementary Fig. 4a). The last source of pathogenic mutations included all disease-causing mutations from UniProtKB. In total, the pathogenic class consisted of 6,713 unique pathogenic mutations (Fig.1 and Supplementary Fig. 2a).

We collected putatively benign variants from the a large population database, gnomAD^34^ (Genome Aggregation Database), and additional population-specific databases, including individuals of Korean^35^ and Japanese^36^ ancestry, as well as human orthologous polymorphisms^37^ (Supplementary Fig. 2a). We defined benign variants as those with a minor allele frequency (MAF) greater than the associated disease prevalence in accordance with ACMG-AMP guidelines^5^. Using a disease-specific MAF threshold, we gained almost 3,000 benign variants compared to using a generic MAF >0.5% threshold (Supplementary Fig. 2b). The benign class consisted of 4,512 variants. The above approach of using gene-specific MAF thresholds can generally be applied to other genes to increase the number of benign variants. Overall, we assembled 40% benign and 60% pathogenic variants (Supplementary Fig. 2d).

### Development of the partners score to quantify epistasis

Overall, DeMAG uses only 13 features, 8 derived from sequence conservation, and 5 from 3D structural models, disorder scores and epistatic relationships (Supplementary Table 4). We designed a novel feature called the partners score, based on the observation that partner residues that are connected whether because they are close in 3D proximity or because they are co-evolving, share the same phenotypic effect (Supplementary Fig. 7a).

We used the AlphaFold2 3D protein structural models to identify residues in spatial proximity (<11Å between C-alpha atoms, see Methods) and highly correlated positions inferred from multiple sequence alignments of homologous sequences^38,39^, to identify co-evolving residue pairs. Each residue position can be associated with only pathogenic, only benign or both pathogenic and benign (mixed), or not being associated with any known mutations (Fig. 2a). Each residue has a score (residue score) based on the type and number of connections it has (Fig. 2a). We used a mixture discriminant analysis^40^ approach to define the partners score: we first estimated the density of the residue score for pathogenic and benign mutations in the training set and then we calculated the posterior probability of belonging in either class, given the residue score. The posterior probability of pathogenicity represents the partners score (Fig. 2a and Methods), which highlights how mutations with the same phenotypic effects cluster both in linear and 3D space of the protein.

**Figure 2:**
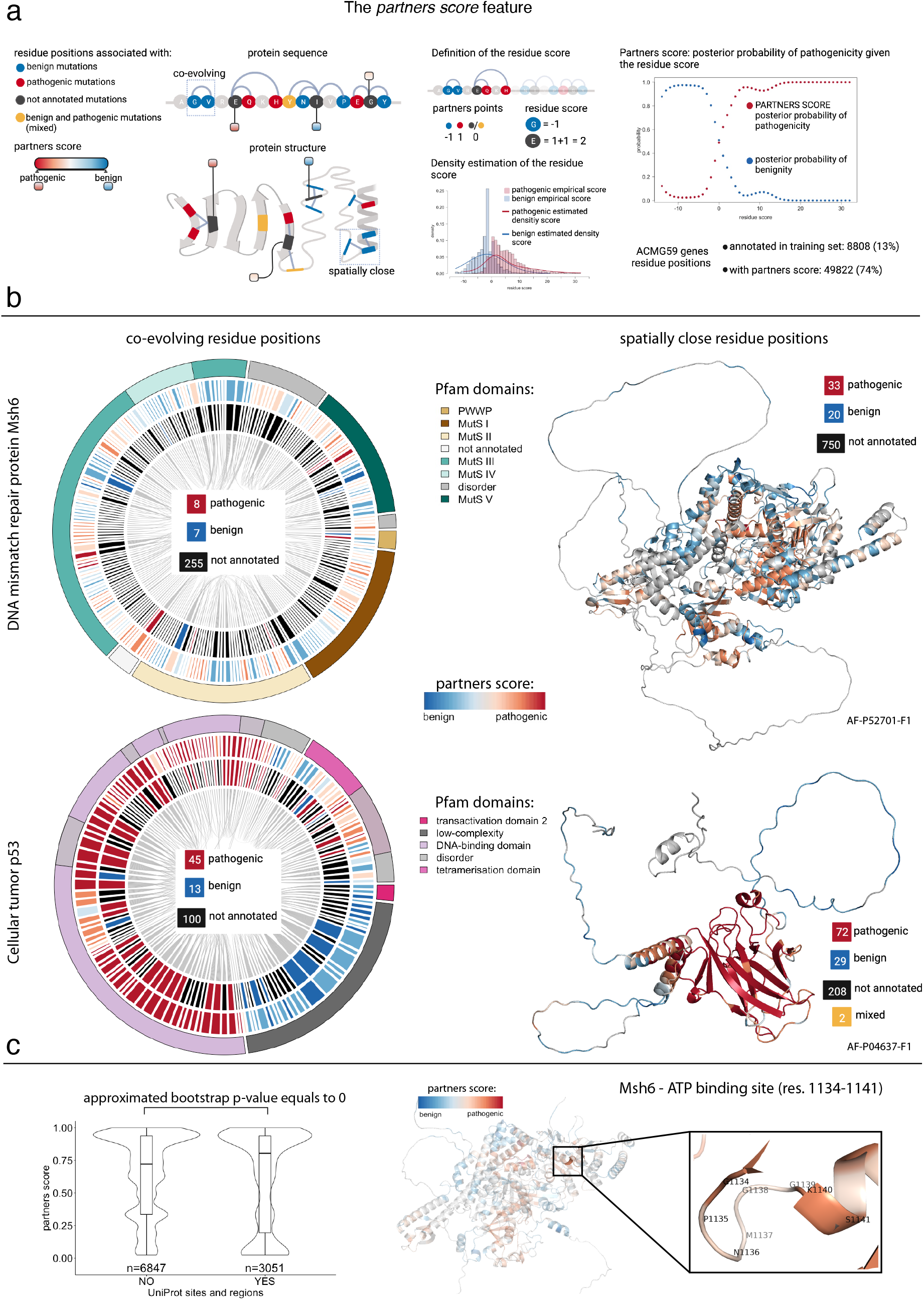
The partners score: integrating evolutionary and structural information to inform variants effect assessment. **a:** The design of the partners score feature. On the left, co-evolving positions in the protein sequence and spatially close residues (<11 Å) in the protein 3D structure. Residues are colored according to the associated mutations’ phenotypic effect, e.g., red for positions with only pathogenic mutations (see legend on the left). Only residues with at least one connection with an annotated residue position have a partners score. On the right, we first defined the residue score as the sum of partners points per residue. Next, we estimated the density of the residue score distribution for the benign and pathogenic residue positions with a gaussian mixture model. Then, for each position we calculated the posterior probability of belonging both to the pathogenic and benign class, given the residue score. The posterior probability of pathogenicity represents the partners score. The partners score feature is available for 74% of ACMG SF residues positions, compared to only 13% of positions annotated in the training set. **b:** On the left, co-evolving residue positions for the DNA mismatch repair protein Msh6. The inner circle shows that there are only 7 pathogenic and 8 benign annotated co-evolving positions. The outer circle shows the partners score, which is also assigned to the 255 not annotated residue positions. On the right, AlphaFold2 3D model of the protein where residue positions are colored based on their partners score. Below, the same representation for the cellular tumor antigen p53. More co-evolving residue positions are annotated and it is clearer a correlation between partners score and domain annotation. **c:** On the left, the partners score distribution between residue positions located (YES) and not found (NO) in important sites e.g., DNA or ATP binding sites. On the right, an example for msh6 protein, whose ATP binding site residues have a high partners score.

### Partners score identifies functional sites

While only 13% of ACMG SF residues positions have annotated mutations in the training set, we can now inform 74% of positions with the partners score by making use of these relationships (Fig. 2a). For example, the DNA mismatch repair protein Msh6 has only 7 pathogenic and 8 benign co-evolving annotated residue positions, but with the partners score we annotated another 255 positions. The same trend applies to positions in spatial proximity (Fig. 2b).

For the cellular tumor protein p53, we observed a clear correlation between the partners score and Pfam protein domain annotations, e.g., the low-complexity region and disordered region are characterized by low partners score, while the DNA-binding domain has overall high score (Fig. 2b).

In addition, we observed that residues located in important functional sites of the protein such as DNA or ATP binding sites, have statistically significantly higher partners score compared to other residue locations (Fig 2c and Methods). Interestingly, we found that the Msh6 ATP binding site has a high partners score (Fig. 2c). The role of the ATP binding site of the Msh2-Msh6 heterodimer is crucial for DNA mismatch repair (MMR) competency: mutations of the lysine invariant residue in the msh6 Walker A motif are complete loss of function mutations *in vivo* in *S. cerevisiae*^41^. Moreover, all 14 mutations (G1134[A,R,E,V], P1135A, N1136D, M1137[T,V], G1138R, G1139[D,C,V], S1141[C,P]) in this site are ClinVar VUSs, hence they do not currently have an interpretation. Overall, 74% of variants that are in motifs and domains are pathogenic in our dataset. In accordance, variants in motifs and domains have higher partners score supporting the efficacy of this feature in assessing the effects of mutations.

### DeMAG reaches high sensitivity and specificity

Many existing VEPs do not explicitly attempt to balance sensitivity and specificity. Their recommended thresholds are usually set to reach high sensitivity in variant interpretation, while tolerating a high misclassification rate of benign variants. This imbalance increases the number of potentially false positive variants to be evaluated (benign variants predicted incorrectly to be pathogenic).

To address this issue, we made efforts to improve training set balance with 60% pathogenic and 40% benign mutations (Fig. 1 and Supplementary Fig. 2d). The 13 features that we selected had independent balanced performance in discriminating between pathogenic and benign classes (Supplementary Table 1a and Methods). DeMAG was trained with a gradient boosting tree method^42,43^ (See Methods) and it yielded high accuracy (87%) and AUC-ROC (92%) values that correspond to high sensitivity (87%) and specificity (85%), as well as high precision (90%) (Fig. 3c). Our recommended threshold to interpret a variant as pathogenic is 0.5. Overall, DeMAG has a balanced sensitivity and specificity that is also maintained within genes (Fig. 3a).

**Figure 3:**
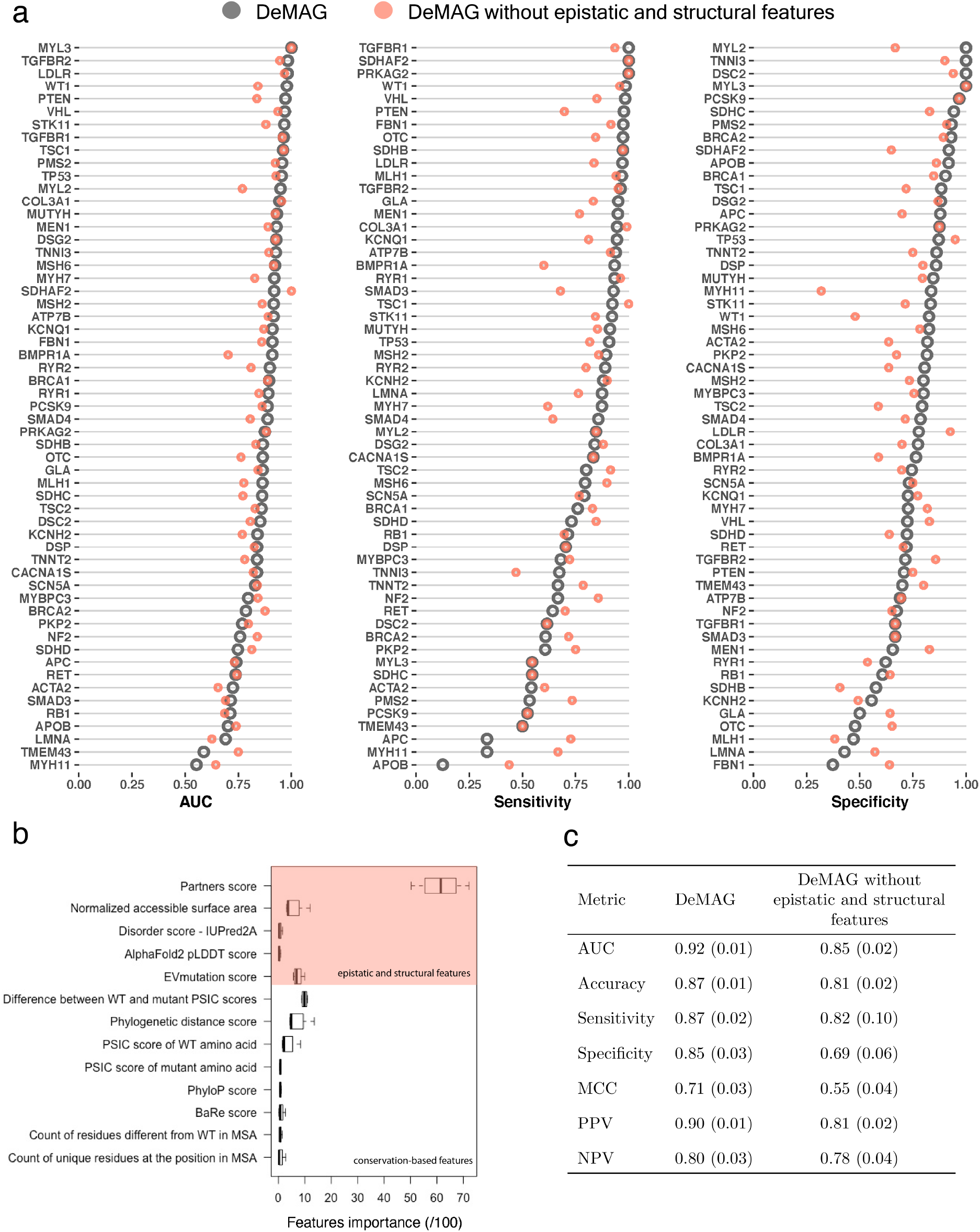
Epistatic and structural features increase DeMAG performance, both within and between genes. **a:** DeMAG performance within genes: for most genes, variants are highly classified (AUC>70%). The high performance is also maintained for the classification of pathogenic (sensitivity>70%) and benign (specificity>70%) variants, regardless of the gene, with only few exceptions. **b:** DeMAG features’ importance for the 4-fold CV. On the x-axis is feature importance and on the y-axis are features names (see Supplemetary Table 4 for features description). The partners score feature clearly stands out from the others. **c:** Different performance metrics for DeMAG. The metrics represent mean values of 4-fold CV with standard deviation in parentheses. Notably, DeMAG is balanced across all evaluation statistics (e.g., 87% sensitivity and 82% specificity). DeMAG without epistatic and structural features has consistently lower performance than the complete model.

### Epistatic and structural information increase DeMAG’s performance

We investigated the contribution of each feature and observed that the partners score is the most informative one (Fig. 3b). In addition, a structural feature, i.e., the normalized accessible surface area, is contributing at least as much as other conservation-based features, e.g., PSIC score. In order to quantify the contribution of epistatic and structural features, we trained DeMAG without those features and observed a consistent decrease across all evaluation metrics (Fig. 3c). In particular, the specificity dropped from 85% to 69%, while the sensitivity was decreased from 87% to 82%.

We explored the contribution of epistatic and structural features at the gene level and observed an increase in sensitivity for genes with high proportions of pathogenic mutations. The specificity increased for genes with different proportions of pathogenic mutations, albeit mainly for genes with high proportions of benign mutations (Supplementary Fig. 8a). The difference in performance with and without epistatic and structural features appears to be independent of the number of training variants per gene (Supplementary Fig. 8b). While it is evident that DeMAG’s performance increased with epistatic and structural features overall, the improvement at the gene level is more challenging to assess.

### Many VEPs fail to predict functional effects observed in deep mutational scans

We validated DeMAG against Deep Mutational Scanning (DMS) data as in prior studies^15^, using data for 4 genes (*BRCA1*^44^, *TP53*^45^, *PTEN*^46^ and *MSH2*^47^), which are expected to be strong proxies for variant functional effects. Most variants in these assays are not yet annotated in ClinVar (51%), and 41% are ClinVar VUSs (Supplementary Fig. 9). Among the 7 VEPs evaluated, DeMAG performed best on *BRCA1* (35% Matthews Correlation Coefficient, MCC) and *PTEN* (27% MCC), while EVE performed better for *TP53* (39% MCC) and *MSH2* (38% MCC) (Supplementary Fig. 9).

Nevertheless, the overall performance of VEPs on functional data is poor, with MCC values below 40% (Supplementary Fig. 9). This might be due to the high proportion of variants interpreted as functional by DMS data, e.g., for *MSH2*, 92% of single nucleotide missense substitutions are assessed as functional. As VEPs usually set recommended thresholds to interpret pathogenicity for high sensitivity, they fail to identify benign variants. While EVE and DeMAG outperform other tools, their misclassification rate is still high on DMS datasets, which underlies a potential limitation for the clinical application of such data.

### DeMAG outperforms existing tools on clinical and population data

As VEPs often collect variants from the same database sources, it is essential to benchmark the different predictors against an unbiased testing set to avoid type 1 circularity^16^. To compare our performance with PolyPhen-2, SIFT4G, REVEL, DEOGEN2^48^, M-CAP, VEST4 and EVE, we designed a clinical testing set comprising high-quality variants submitted to ClinVar after 2017. As the most recent supervised method was published in 2016, none of the VEPs were trained on those newer variants. We re-trained DeMAG without the testing variants and used this model to predict ClinVar testing variants (Supplementary Table 2). The testing set had 852 (66%) pathogenic and 433 (34%) benign variants. As not all VEPs have predictions for these variants, we benchmarked DeMAG in pairs (Table 1 and Fig. 4a). DeMAG stood out from all the other predictors, reaching the highest specificity, accuracy, and MCC value (Table 1 and Fig. 4a). DeMAG performance is consistently high across the different evaluation metrics. There is great discrepancy between the specificity of DeMAG and other tools, e.g., DeMAG 84% and REVEL 62%. It should be noted that EVE’s high accuracy (89%) dropped to 72% when we included variants that the authors predicted as *Uncertain*, although they are actually annotated as benign or pathogenic variants in ClinVar. While each tool has different strengths, DeMAG outperforms all other methods tested in at least one evaluation metric reported.

**Table 1.** DeMAG best classifies clinical variants. Different comparison metrics for DeMAG and seven popular pathogenicity predictors tools. The test set is assembled from the ClinVar database, consisting of both pathogenic (n=852) and benign variants (n=433) submitted after the year 2017. The comparison in pairs guarantees that each predictor is evaluated on all the variants for which a prediction exists. DeMAG is the best performing tool, as it is the most balanced across all of the metrics. DeMAG’s specificity highlights how it does not overpredict pathogenic variants like other tools such as SIFT4G and M-CAP.

| VEPs | Sensitivity | Specificity | Accuracy | MCC | AUC | Variants Predicted (n = 1285) | VEP's coverage |
| --- | --- | --- | --- | --- | --- | --- | --- |
| DeMAG | 0.93 | 0.82 | 0.9 | 0.77 | 0.96 |  | 0.95 |
| SIFT4G | 1 | 0.07 | 0.68 | 0.21 | 0.81 | 1226 | 0.95 |
| DeMAG | 0.94 | 0.82 | 0.89 | 0.76 | 0.96 |  | 1 |
| REVEL | 0.99 | 0.62 | 0.86 | 0.69 | 0.96 | 1285 | 1 |
| DeMAG | 0.93 | 0.79 | 0.89 | 0.74 | 0.95 |  | 0.75 |
| DEOGEN2 | 0.91 | 0.58 | 0.8 | 0.53 | 0.91 | 964 | 0.75 |
| DeMAG | 0.93 | 0.79 | 0.89 | 0.74 | 0.95 |  | 0.76 |
| PolyPhen2 | 0.91 | 0.72 | 0.85 | 0.64 | 0.86 | 972 | 0.76 |
| DeMAG | 0.94 | 0.81 | 0.89 | 0.76 | 0.96 |  | 1 |
| VEST4 | 0.99 | 0.64 | 0.87 | 0.72 | 0.95 | 1280 | 1 |
| DeMAG | 0.94 | 0.81 | 0.9 | 0.76 | 0.96 |  | 0.98 |
| M-CAP | 1 | 0.28 | 0.77 | 0.44 | 0.93 | 1256 | 0.98 |
| DeMAG | 0.96 | 0.81 | 0.92 | 0.8 | 0.97 |  | 0.73 |
| EVE | 0.92 | 0.81 | 0.89/0.72<sup>1</sup> | 0.73 | 0.91 | 940 | 0.73 |

<sup>1</sup>The accuracy is 72% if misclassified variants based on EVE's Uncertain class are included.

**Figure 4:**
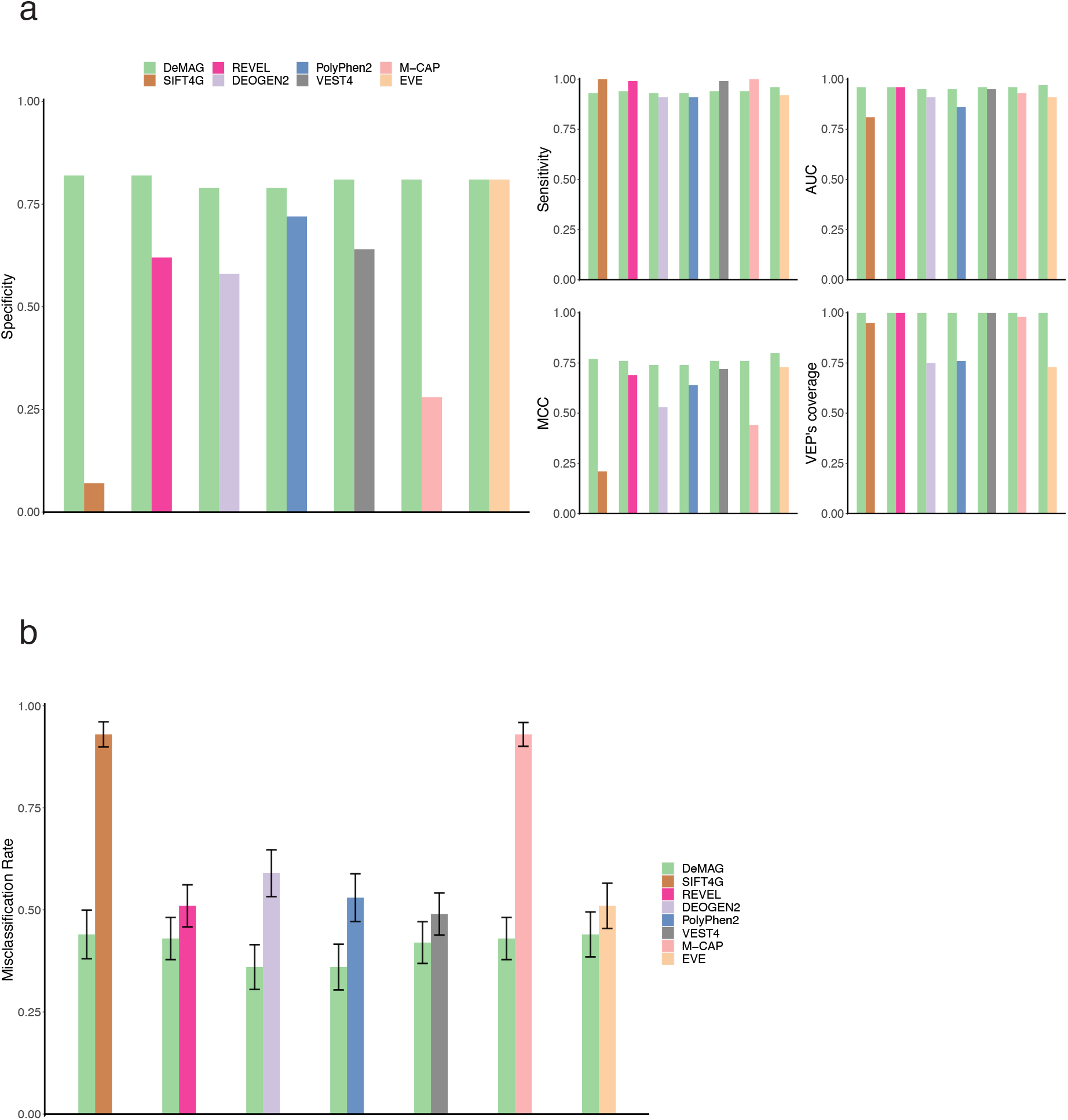
DeMAG outperforms VEPs on clinical and putatively benign common variants. **a:** DeMAG’s pairwise comparison on clinical variants from the ClinVar testing set, according to difference performance metrics (specificity, sensitivity, ROC-AUC, Matthews Correlation Coefficient (MCC), variant coverage). The left panel shows that, except for EVE, DeMAG’s advancement in specificity stands out. While all tools reach almost perfect ROC-AUC and sensitivity (top right panels), DeMAG reaches the most balanced performance, namely the highest MCC (bottom left panel). **b:** DeMAG’s pairwise comparison on putatively benign common variants from the Estonian Biobank. DeMAG shows the lowest misclassification rate among all VEPs.

We also benchmarked against putatively benign common variants in the Estonian population^49^. Even if variants from the Estonian Biobank are not yet part of gnomAD, 80% were already annotated in ClinVar (Table 2 and Fig. 4b). Most variants were VUSs (33%) and high-quality benign variants (30%). We evaluated variants, not already annotated in ClinVar or in our training set, and we filtered the variants based on MAF greater than the corresponding disease prevalence, resulting in a total of 94 putatively benign variants. DeMAG had by far the lowest misclassification rate among the other predictors (Table 2 and Fig. 4b). DeMAG yielded superior performance across clinical and population-based validation sets.

**Table 2.**
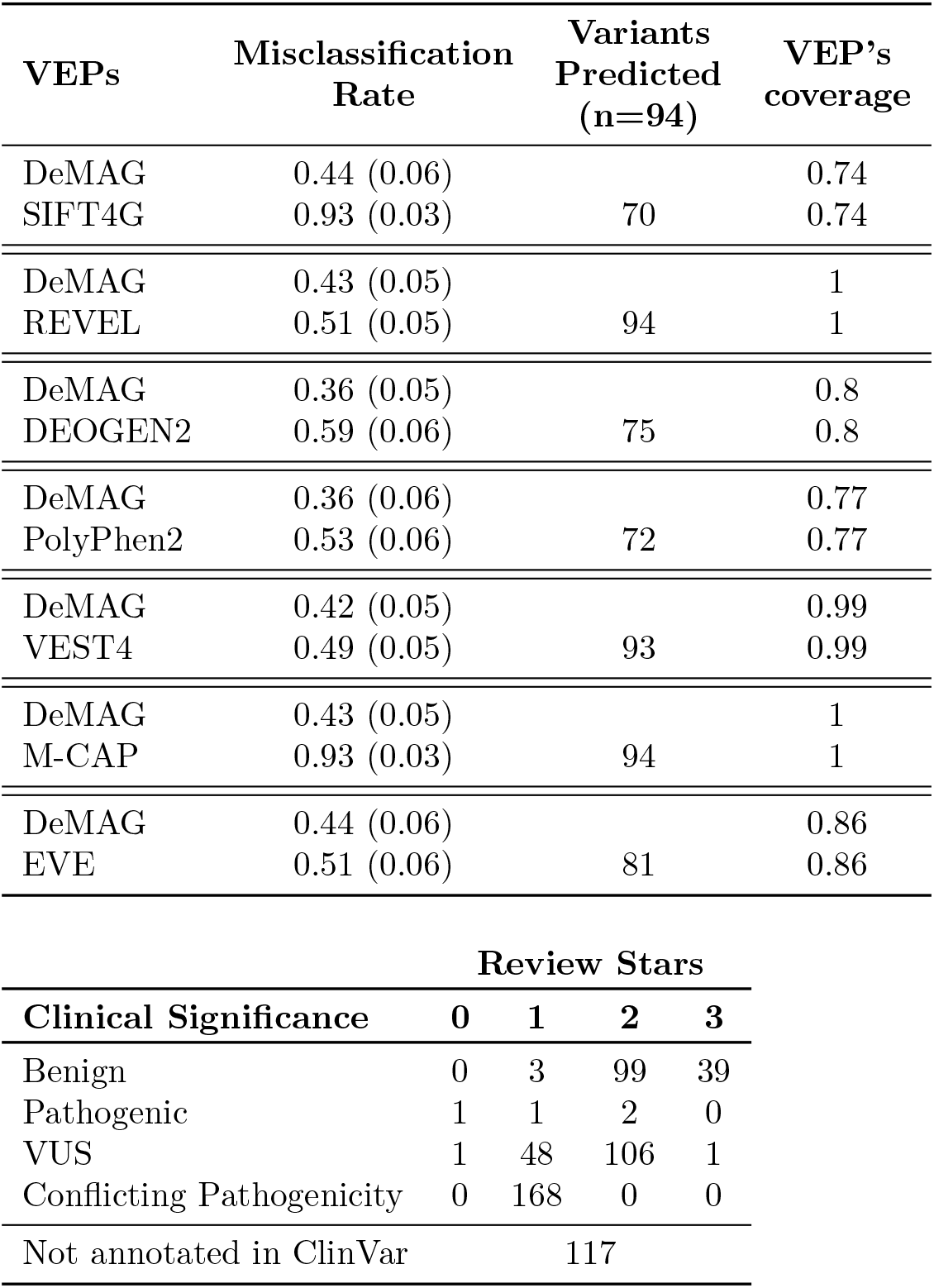
DeMAG classifies putatively benign variants with the lowest misclassification rate. Misclassification rate of putatively benign common variants within the Estonian population (see Methods). DeMAG reaches by far the lowest misclassification rate among the other tools. The values in parentheses represent the misclassification rate standard deviation calculated on 5000 bootstrap samples. The lower table shows ClinVar clinical significance for the Estonian Biobank variants: among high-quality (review status with at least 2 stars) annotations most variants are benign and VUSs.

## Discussion

As genomic sequencing becomes more frequent in clinical practice and research, the interpretation of missense variants remains a major challenge. It is especially important to correctly classify genetic variants in the context of clinical decision making, i.e., variants in clinically actionable genes. Therefore, we developed DeMAG, a specialized VEP tool that reaches high performance for clinically actionable disease genes, as defined by the ACMG. It demonstrates superior balance between sensitivity and specificity (Fig. 3a and c), and can be useful for variant prioritization, rare variant studies, and to reclassify VUSs in the ACMG SF genes. As high as 30% of missense VUSs in ClinVar belong to the ACMG SF genes, underlying the importance of a specialized classifier. While exome-wide predictors have a wide range of utilities in basic research, here we show that a specialized classifier reaches higher performance on clinically actionable genes, and should be prioritized in translational research (Tables 1 and 2 and Fig. 4).

The assembly of a high-confidence balanced training set is crucial for the development of supervised predictors. For example, the ClinVar *Review status* provides a system to evaluate review quality and agreement on the clinical significance of a variant. Thus, we included only variants whose clinical interpretation is shared among different submitters, i.e., 2 or more review status stars. On the other hand, HGMD does not provide detailed rules to distinguish between high and low-quality variants’ annotations. The joint analysis of clinical annotations between databases, namely ClinVar and HGMD, allowed the removal of potentially conflicting or lower-quality variants. Indeed, we removed almost 40% of disease-causing variants in HGMD that were interpreted in ClinVar as VUSs (Supplementary Fig. 4). As clinical databases become increasingly important repositories for genetic variation in relation to human health and disease phenotypes, it is crucial to implement quality control pipelines to include only variants with non-conflicting and clear interpretations.

As many VUSs are identified in diagnostic testing, many studies are focusing on VUS assessment and reclassification^50,51^. For instance, *Dines* et al. reclassified *BRCA1* exon 11 as a coldspot, suggesting a strong benign reinterpretation of variants located within that region. DeMAG predictions for that region agree with such reclassification (Supplementary Fig. 11). On the other hand, the reassignment of the *BRCA1* coiled-coil domain (1393-1424) as a moderate benign region seem in disagreement with previous study that showed that mutations in that region disrupt the complex formation with *PALB2*, which would impair the Homologous Repair (HR) mechanism^52^. DeMAG agrees with this work and it classifies at least 45% of all possible missense substitutions in that region as pathogenic (Supplementary Fig. 11). Overall, we provide classification for all missense variants, including all VUSs, revealing that 43% of them are predicted pathogenic by DeMAG.

It is still not common practice to report different performance statistics metrics for VEPs, and often only the AUC-ROC is provided, that is not adequate especially when training on unbalanced data^53^ (Supplementary Fig. 2d). Therefore, we reported several performance metrics when benchmarking DeMAG (Tables 1 and 2 and Figure 4) that highlighted how several popular VEPs fail to identify benign variants (Tables 1 and 2 and Fig. 4). As computational evidence is one of the criteria used to classify sequence variants, bias to overestimate pathogenicity contributes to labeling more variants as pathogenic in publicly available databases.

So far, the lack of experimental structures for most of the human proteome failed to highlight the role of structural properties of individual proteins for most VEPs. For the ACMG SF genes the experimental structures with a resolution higher than 4 Å covered only 28% of the residues (Supplementary Fig. 5), and now all residues have AlphaFold2 3D model predictions. Nevertheless, the comparison between AlphaFold2 and IUPred2A showed how high-quality predictions are still missing for many ordered regions (Supplementary Fig. 6), highlighting how the mystery of protein folding is yet to be understood.

The epistatic and structurally derived features are informative, as DeMAG has inferior performance without these features in training for all metrics considered (Fig. 3c). Despite the overall improvement, there are a few genes that do not benefit from those new features. This might be due to the imbalanced nature of pathogenic and benign training variants within the genes (Supplementary Fig. 2). The performance of genes that harbor almost only pathogenic (or benign) mutations will be dominated by high sensitivity (or specificity), so an improvement in the dominant metric will result in a substantial drop of the other one. For instance, the new features increase sensitivity in FBN1, but as the gene has 93% pathogenic mutations, the specificity drops by 27% (Supplementary Fig. 12 and Fig. 3a). The same happens for APOB, MYH11 and APC. They have benign mutations in proportions greater than 86%, and indeed, an increase in specificity corresponds to more than 30% drop in sensitivity. Though we are able to dramatically improve the balance in performance characteristics, some clinically actionable disease genes have significant biases which pose challenges for variant interpretation.

We combined co-evolutionary and structural information with the annotated phenotypic effect of the coupled positions. We observed that evolutionary coupled positions as well as spatially proximal ones are enriched for the same phenotypic effect and might serve to identify functional sites, e.g., binding sites (Fig. 2c and Supplementary Fig. 7a). We show that the traditional conservation paradigm to interpret human coding missense mutations should be complemented with epistatic and structural information.

DeMAG requires abundant clinical data to be successfully extended to any other gene, which is currently not available for most genes, even among actionable ones. In the coming years, the lack of clinical data for less studied genes will be generated and become available and supervised predictors like DeMAG will further improve variant effect assessment directly from patients sequencing data.

In conclusion, we anticipate that our tool and the web server (demag.org) will facilitate clinical decision-making. Moreover, the newly developed features can be applied to other genotype-phenotype predictors and be generalized to other genes and organisms.

## Acknowledgments

This work was supported by NIH/NHGRI grant (R01-HG010372) and by Max Planck Society MPRGL funding. We thank the Institute of Genomics of the University of Tartu for sharing genotype data from 4,776 individuals. We especially thank Prof. Dr. Andres Metspalu and Krista-Roberta Saviauk for the Estonian Biobank data. We thank Shamil Sunyaev for many scientific discussions and for the critical reading of the manuscript. We further thank Michele Marass and Jonas Pöhls for providing valuable feedback on the manuscript, HongKee Moon for the technical support, and the MPI-CBG Computer Services and Scientific Computing Facility for their support.

## Online methods

### Training dataset

VCF files for SNPs were collected both from clinical and population databases. We downloaded the ClinVar VCF file, version 2021.05, and we retained variants with at least 2 review status stars variants annotations that contained in the clinical significance field either the word ‘pathogenic’ or ‘benign’. Variants of conflicting pathogenicity were excluded as well as variants with only somatic labels. We used the Human Mutation Gene Database (HGMD), version 2020.03, to extract pathogenic mutations. We filtered for disease mutations (DMs) and we only retained variants that were not already annotated in ClinVar (Supplementary Fig. 4). With this analysis we removed HGMD variants with a VUSs label in ClinVar (26%), as well as low quality (zero and one review status star) ClinVar pathogenic variants (27%) and ClinVar benign variants (1%). PolyPhen-2 HumVar set derived from UniProtKB release 2021_01 was utilized to collect both pathogenic and benign variants. We also added common variants to the benign set from the Genome Aggregation Database (gnomAD), from the NCBI ALFA (Allele Frequency Aggregator) project release 20201124, from country specific sequencing projects, i.e., Korea (KRGDB) and Japan (3.5KJPNv2), and from human orthologues variations (PrimateAI and HumOrtho). For the population data, as suggested by the ACMG-AMP guidelines, we considered as benign the variants with a minor allele frequency (MAF) greater than the associated disease prevalence. Disease prevalences were collected from Orphanet and when not available a MAF greater than 0.5% was applied (Supplementary Table 3). Putatively benign non-human primate variants were collected from primateAI database but only Chimpanzee and Bonobo species were considered. Both duplicates and conflicting variants, i.e., variants reported both as pathogenic and benign among different sources, were removed from the final training set. Number of training variants among different sources and different genes are shown in Supplementary Fig. 2.

### Clinical testing dataset

The primary clinical testing set (852 pathogenic and 433 benign variants) was built from the ClinVar database. To ensure the independence of the testing set, we only considered variants submitted to ClinVar after December 2017 (Supplementary Fig. 3). Since the newest supervised method we were benchmarking with was published in 2016, we are sure that those variants were not used in the training pipeline of any of these predictors. As we used ClinVar variants for training, we trained a different model excluding the variants used as a testing set. This ensured unbiased comparison for DeMAG (Supplementary Table 2).

### Functional variants testing set

In order to investigate the accordance between DeMAG predictions and experimental data, we evaluated DMS data for *BRCA1, TP53, PTEN* and *MSH2*. All datasets were collected from the Supplementary material of the respective papers. When possible, we assessed the concordance between different experimental replicas to ensure a robust functional score for each variant. For *BRCA1*, two scores were available and as the correlation and the variance explained was 81% and 65% respectively, we included all the variants. The authors assessed variants’ functional scores in three categories: loss of function (LOF), intermediate (INT) and functional (FUN). We did not evaluate the intermediate class. After removing overlapping variants with our training set, we evaluated 1587 variants: 1268 (80%) FUNC and 319 (20%) LOF. For *PTEN*, 8 different scores were available. Since the correlation pattern among the replica was variable, we only evaluated variants whose standard deviation among all available 8 scores was smaller than 10%. In this case as well, we did not evaluate uncertain functional categories, namely *possibly wt-like* and *possibly low*. We eventually evaluated 34 FUNC (64%) and 19 LOF (36%) variants. For *MSH2*, we could not analyze the concordance among different replicas, as only one score was provided. The total number of variants analyzed was 5075: 4737 (93%) FUNC and 338 (7%) LOF variants. The last gene we analyzed was *TP53*, for which we did not have more than one functional score but agreement between replicas was already assessed by the authors. We eventually evaluated 1017 variants: 714 (70%) FUNC and 303 (30%) LOF variants.

### Common variant testing set

We assembled another testing set from the Estonian Biobank. This set comprises common variants in the Estonian population. To consider a variant as benign we applied the same rule as for the common variants in the training set: MAF greater than disease prevalence. In order to design a testing set as independent as possible, we removed variants that were present in our training set as well as variants with a ClinVar interpretation. This might not guarantee that those variants were not used for training by the other tools, thus there is still a potential bias to overestimate their performance. The validation set consisted of 94 variants. We repeated the analysis on 5000 bootstrap samples, in order to obtain the standard deviation of the performance metric, namely the misclassification rate.

### Pathogenicity scores

Pathogenicity scores were collected through dbNSFP v4.1a command-line application. We have downloaded scores for SIFT4G v2.4, VEST v4.0, Polyphen-2 v2.2.2, M-CAP v1.3, DEOGEN2 and REVEL. To calculate the accuracy, we used the threshold as recommended by the authors, which is 0.5 for all the methods except for M-CAP which is 0.025 and SIFT4G which is 0.05. For EVE, we downloaded the predictions from the web server (https://evemodel.org/download/bulk). EVE does not provide a unique threshold, rather a gene-based predefined categorical feature with 3 different levels: Pathogenic, Benign and Uncertain. We evaluated only the benign and pathogenic variants, but we calculated the accuracy when misclassified variants based on EVE’s *Uncertain* class were retained.

### Variant annotation

We used PolyPhen-2 MapSNPs annotation tool to map the genome assembly hg19/GRCh37 variants coordinates to missense coding SNPs. Only variants mapping to known canonical transcripts according to the UCSC Genome Browser were retained.

### Sequence-based features

We used PolyPhen-2 pipeline to annotate DeMAG features, a complete list and description is available at the PolyPhen-2 Wiki page (http://genetics.bwh.harvard.edu/wiki/pph2/appendix_a). The new features are annotated separately (see sections below). IUpred2A scores were collected from the tool’s web interface (https://iupred2a.elte.hu/).

### Epistatic and structure-based features

EVmutation scores were obtained using the EVcouplings python package, version v0.1.1. The alignments were manually curated in order to maximize the coverage. In particular, protein sequences were tiled in regions of 100 residues with an overlapping window of 50 residues, i.e., 1-100, 50-150. We computed the first three stages of the EVcouplings pipeline, i.e., align, couplings and compare, for each tiled region and for five different bit score thresholds (0.1,0.2,0.3,0.4,0.5). We merged adjacent regions if either the number of sequences in the alignment was greater than 5 times the length of the region or if the skewness of the Evolutionary Couplings (EC) distribution was greater than 1. We repeated these steps until no more adjacent regions could be joined together. The final alignment coverage for the ACMG SF genes is shown in Supplementary Fig. 12a. Co-evolving residue positions were identified within the EVcouplings framework. Each residue might be associated with many or only few residue positions.

We excluded residue positions in a connection with only not annotated residues as in the training set. A residue that is not annotated is a residue that according to our training set does not have any associated mutations. To each residue is assigned a score that is 1 for pathogenic residue positions, −1 for benign and 0 for mixed or not annotated ones. The residue score is the sum of the scores of all co-evolving positions (Fig. 2a). We used the residue score distribution as input for the mixture discriminant analysis. First, we estimated the density of the residue score distribution for the pathogenic and benign residues positions independently. Next, we predicted for each residue position the posterior probability of belonging to the benign and pathogenic class given the residue score and the prior probability of being a pathogenic or benign position as in the training set. The partners score of co-evolving residue positions is the posterior probability of belonging to the pathogenic class (Fig. 2a).

The significance of evolutionary coupled residues is determined by their location in the EC distribution. A probability model has been defined to identify strong coupled positions. The higher the probability the stronger the coupled residues. In order to select the best probability threshold, we trained such a model for different cutoffs. We selected the probability cutoff (0.6) with the smallest difference between sensitivity and specificity (Supplementary Fig. 7b). The same approach was used for spatially close residue positions as in AlphaFold2 3D models. In order to select the Ångström distance threshold for considering a pair of residues as contacting in 3D space we trained different models with different cutoffs (4-11Å). We selected 11Å as the best distance, i.e., smallest difference between sensitivity and specificity (Supplementary Fig. 7b).

We did not consider larger distances to avoid introducing protein-specific properties rather than residues-based ones.

The residue score for co-evolving positions and spatially close ones highly correlate (Supplementary Fig. 7c), thus we combined them and in case of overlap we took the spatially close residues score. To derive the partners score feature we used a mixture discriminant analysis approach implemented with the mclust^54,55^package in R. The best model selected by BIC had 3 gaussian components with variable variance for the density of the residue score for pathogenic residue positions and 4 gaussian components with equal variance for the benign ones. Given the density estimation of the residue score and the prior probabilities of the pathogenic and benign residue positions in the training set, the mixture model predicts the posterior probability of belonging in both classes (pathogenic and benign). We considered the posterior probability of pathogenicity as the partners score. Eventually, 49822 residues have a partners score. Both the residue score and the partners score are not biased towards protein’s length, thus scores normalization was not needed (Supplementary Fig. 7c).

In order to evaluate whether mutations located in functional motifs and domains have higher partners score than any other locations, we tested the null hypothesis of equal means in the two groups with a bootstrap analysis approach. We first built the distribution under the null hypothesis of equal means in the two groups. Then, we drew 20000 samples from the null distribution and stored the t-statistic of each sample. Lastly, we calculated the approximated p-value by dividing the times the t-statistic in each of the bootstrap samples was greater than t-statistic observed in the original sample by the number of bootstrap samples. The approximated p-value was 0. We obtained the same statistically significant results with the Wilcoxon rank sum test (p-value was 0.0005532).

### Structures and 3D models

Structures were collected from PDB (Protein Data Bank). We included any experimental structure with a resolution not greater than 4 Ångström. To increase structural coverage, we also included 3D models. We used SWISS-MODEL (https://swissmodel.expasy.org/) to derive protein structure homology models, as well as *de novo* modeling. For the homology modeling we only considered models with a template identity greater than 70% to the query protein. We used the EVcouplings pipeline as already described in the Features paragraph. For each protein’s region we got different 3D models predictions. The models were clustered based on their RMSD score. The most populous cluster should contain the most reliable models. To select the best model, we chose the top 20 models that belonged to the biggest cluster. Among those we excluded models with knots and we took the model with the smallest radius of gyration, as a metric of protein compactness^56^. Overall, considering those 3 different sources for 3D structures/models, 50% of ACMG SF residues were covered (Supplementary Figure 5a). Since July 2021 we updated our 3D data with AlphaFold2 models (https://ftp.ebi.ac.uk/pub/databases/alphafold/UP000005640_9606_HUMAN.tar) that resulted in 100% coverage among ACMG SF genes (Supplementary Fig. 5). For long genes (APC, APOB, BRCA2, DSP, FBN1, RYR1, RYR2) AlphaFold2 produces different overlapping models that we combined to obtain one single complete model. The models are ∼1400 aa long with non-overlapping regions of ∼200 aa that eventually cover the full sequence.

### Cross-validation scheme

In order to select the probability and Ångström distance cutoff as well as for the features selection pipeline and in general for the model training, we trained the models with a cross validation scheme. The cross-validation scheme ensured that each testing fold contained different proteins than the ones in the training folds. This prevents bias due to training and testing within the same protein. In addition, each testing fold should have a distribution of the pathogenic and benign class that reflects the one of the training set. To respect these two principles, we considered 4 CV-folds for model training and hyperparameters selection and 5 CV-folds for the features selection pipeline.

### Feature selection

Training set variants were annotated with a total of 91 features. Most features were annotated with PolyPhen-2 pipeline and a description of each feature can be found here (http://genetics.bwh.harvard.edu/wiki/pph2/appendix_a). After removing gene-based features, 39 features were retained. In order to select the most discriminative features to train the model with, we trained a univariate logistic regression model. The CV strategy is explained above (see Cross-validation scheme subsection). The features that had an ROC-AUC above 0.7, while ensuring a correspondent sensitivity and specificity greater than 0.5, were selected. We trained the model with a total of 13 features. The feature selection process was repeated twice: one time for DeMAG and another one for DeMAG without the ClinVar testing set. The selected features for both models can be found in Supplementary Tables 1a and 2b.

### Machine learning models

DeMAG was trained with a gradient-boosting model with classification tree as base learner and Bernoulli deviance as loss function. R package “gbm” version 2.1.8 was used for training^57^. We trained the model with the 13 features selected during the feature selection pipeline. We implemented a grid search for two of the parameters of the gbm function: shrinkage and interaction depth. The combinations evaluated were 9, resulting from 3 values for the shrinkage parameter (0.001, 0.0055, 0.01) and for the interaction depth (1, 2, 3). The best combination of parameters was selected based on performance in 4-fold CV: the models were ranked based on the smallest difference between sensitivity and specificity and if more than one model satisfied the condition the model with the highest sensitivity and specificity was selected (Supplementary Tables 1b and 2b). As for the feature selection, the grid search was performed for DeMAG without the ClinVar test set as well. Once we identified the best parameters, the gbm model was trained with 4-fold CV to inspect the robustness of the 4 models’ performance and to investigate any potential biases in the training set. The final model was then trained on the complete dataset. The gradient boosting model was chosen over other ensemble machine learning techniques such as Random Forest because it explicitly handles missing values, namely for each decision in the tree there are not only the left and right nodes but a missing node as well. Missing information is thus treated as information and it is not imputed.

## Data availability (code and website about training data and precomputed scores)

The code will be available here https://github.com/Fedeluppi/DeMAG and the webserver is https://demag.org/.

## Software

The statistical analysis was all done in R (see the github repository for the lists of packages used). The figures and tables were made with R, Biorender.com and LaTeX. To visualize protein 3D models we used <u>Pymol</u>. The webserver was created with RShiny app.

## List of Supplementary Figures and Tables

Supplementary Figure 1: ClinVar clinical significance is biased for actionable genes.

Supplementary Figure 2: DeMAG training set reaches high balance between the pathogenic and benign class both between and within genes.

Supplementary Figure 3: ClinVar unbiased testing set allows to evaluate 853 pathogenic and 433 variants.

Supplementary Figure 4: Most HGMD variants have a ClinVar label.

Supplementary Figure 5: AlphaFold2 predictions cover 100% of ACMG SF sites.

Supplementary Figure 6: High correlation between IUPred2A disorder and AlphaFold2 confidence score.

Supplementary Figure 7: Spatially close and co-evolving residues are enriched in the same phenotypic effect.

Supplementary Figure 8: Epistatic and structural features increase sensitivity for “pathogenic” genes and specificity for “benign” genes.

Supplementary Figure 9: Most ClinVar VUSs are predicted as functional variants by DMS data.

Supplementary Figure 10: EVmutation covers 73% of ACMG SF sites and 23% of residues are disordered.

Supplementary Figure 11: Interaction with *PALB2* region (1397-1424) in *BRCA1* is mainly predicted as pathogenic by DeMAG.

Supplementary Figure 12: Most genes benefit from the epistatic and structural features.

Supplementary Table 1: DeMAG model is balanced among different statistics metrics.

Supplementary Table 2: DeMAG (ClinVar) model is balanced among different statistics metrics.

Supplementary Table 3: ACMG SF genes disease prevalences (SupplementaryTable3.xlsx)

Supplementary Table 4: DeMAG features

